# NKp46 Recognizes the Hyphal Form of Candida albicans and Mediates Protective Antifungal Immunity

**DOI:** 10.1101/2025.05.02.651874

**Authors:** Mingdong Liu, Ahmed Rishiq, Yoav Charpak-Amikam, Fubin Li, Ofer Mandelboim

## Abstract

*Candida albicans* is an opportunistic fungal pathogen capable of transitioning between yeast and hyphal forms, a morphological plasticity critical for its pathogenicity. Natural killer (NK) cells play a crucial role in antifungal immunity, yet the molecular basis of their interaction with *C. albicans* remains incompletely understood. Here, we identify NKp46 (NCR1 in mice), an activating receptor on NK cells, as a functional receptor for the hyphal—but not yeast—form of *C. albicans*. Using image flow cytometry and binding assays with NKp46-Ig and NCR1-Ig fusion proteins, we demonstrate that NKp46 selectively binds to hyphae. This interaction is mediated by the D2 domain of NKp46 and is sialic acid independent. Blocking NKp46 impairs NK cell degranulation and fungal killing in vitro. In vivo, mice deficient in NCR1 exhibit increased susceptibility to systemic *C. albicans* infection. Our findings establish NKp46 as a key sensor of invasive fungal morphology and underscore its role in early antifungal immunity.

## Introduction

*Candida albicans* is a commensal fungus that asymptomatically colonizes human mucosal surfaces such as the skin, oral cavity, gastrointestinal tract, and urogenital tract[1]. However, under conditions of immune suppression or antibiotic use, *C. albicans* can transition from a benign commensal to a pathogenic invader, leading to mucosal infections or life-threatening systemic candidiasis[2]. Despite antifungal therapy, mortality from systemic *C. albicans* infections often exceeds 40%[3,4].

One of the major virulence traits of *C. albicans* is its ability to undergo morphological transitions between yeast, pseudohyphal, and true hyphal forms[2]. The yeast form is associated with colonization and dissemination, whereas the hyphal form exhibits enhanced tissue penetration and immune evasion[3]. Recognition and clearance of the hyphal form are thus essential components of effective antifungal immunity[5,6].

Natural killer (NK) cells are innate lymphocytes that contribute to host defense against tumors, viruses, and fungi[7–11]. Their cytotoxic activity is regulated by a balance of activating and inhibitory receptors [12]. In fungal immunity, NK cells have been shown to utilize receptors such as TIGIT and NKG2D to recognize *C. albicans*, while NKp46 is implicated in the recognition of *C. glabrata*, but not *C. albicans*[13– 15].

NKp46 (NCR1 in mice) is a natural cytotoxicity receptor that mediates activation of NK cells in response to a wide array of pathogens [14,16]. It contains two extracellular immunoglobulin-like domains (D1 and D2), with the D2 domain playing a central role in ligand binding[17]. Here, we provide evidence that NKp46 does bind *C. albicans*—specifically in its hyphal form—thus redefining the role of NKp46 in fungal immunity.

## Results

### NKp46-Ig and NCR1-Ig specifically recognize the hyphal form of *C. albicans*

To investigate the potential recognition of *C. albicans* by NKp46, we stained *C. albicans* in different morphological states. *C. albicans* cultured at 30°C in Sabouraud medium remained in the yeast form, while incubation in RPMI at 37°C induced hyphal formation. Cells were stained with NKp46-Ig or NCR1-Ig and analyzed by flow cytometry. Gating based on forward and side scatter parameters allowed separation of yeast and hyphal populations (Figure 1A, left). We observed strong binding of both NKp46-Ig and NCR1-Ig to the hyphal form, while no significant binding was detected on the yeast form (Figure 1A, right).

**Figure 1.**
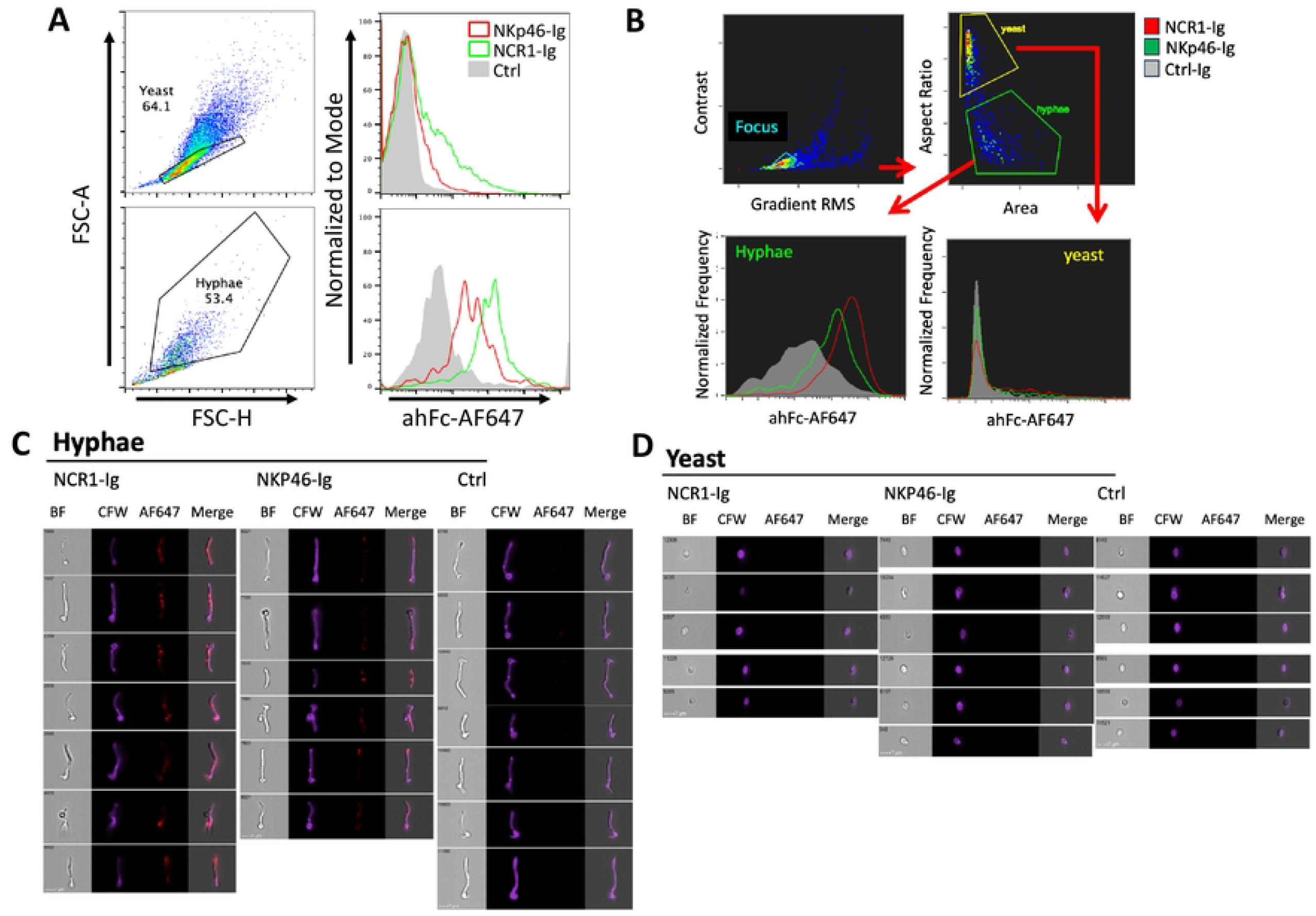
NKp46-Ig and NCR1-Ig specifically bind to the hyphal form of *C. albicans*. (A) *C. albicans* was cultured in Sabouraud medium at 30°C to maintain the yeast form (upper left) or in RPMI at 37°C (lower left) to induce hyphal formation. Cells were stained with NKp46-Ig or NCR1-Ig (right histograms), followed by an anti-human Fc Alexa Fluor 647 (AF647)-conjugated secondary antibody. Experiments were repeated three times. (B) Gating strategy for image flow cytometry analysis of *C. albicans* cultured in RPMI at 37°C. Cells were stained with Calcofluor White (CFW) and NKp46-Ig or NCR1-Ig, followed by ahFc-AF647. Focused cells were initially gated based on “Contrast” and “Gradient RMS.” Yeast and hyphal forms were then distinguished using “Aspect Ratio” and “Area.” (C) Representative images of *C. albicans* in the yeast form from (B). Brightfield (BF), CFW staining, fusion protein staining, and the merged channel are shown. (D) Representative images of *C. albicans* in the hyphal form from (B). Brightfield (BF), CFW staining, fusion protein staining, and the merged channel are shown.

To determine whether NKp46 binding is driven by morphological changes or culture conditions, we further analyzed *C. albicans* cultured in RPMI at 37°C for 3 hours using imaging flow cytometry. Morphological distinction between yeast and hyphae was achieved by gating focused cells based on “Aspect Ratio” and “Area” parameters (Figure 1B, upper). Tightly gated populations of *C. albicans* yeast and hyphal forms were compared by immunofluorescence intensity to evaluate the differential binding of fusion proteins to the two fungal morphologies. We observed that significant binding of NCR1-Ig and NKp46-Ig occurred only in the hyphal form, whereas minimal or no binding was detected in the yeast form (Figure 1B, lower).

To further validate the accuracy of yeast and hyphae gating, representative images of stained hyphal and yeast form cells were presented (Figure 1C-D). Consistent with our previous results, NKp46-Ig and NCR1-Ig bound almost exclusively to hyphal cells. The binding of NKp46-Ig and NCR1-Ig was not restricted to the elongated hyphal segments but was also observed on the yeast-like regions of the fungal cells. These findings indicate that the morphological switch to hyphae, rather than the culture environment itself, determines NKp46 ligand expression.

### Structural requirements and binding specificity of NKp46 to *C. albicans*

To dissect the structural basis of NKp46 binding to hyphal *C. albicans*, we used fusion proteins containing individual NKp46 domains (D1-Ig and D2-Ig). Flow cytometry revealed that the D2 domain alone was sufficient to mediate binding to hyphal *C. albicans*, indicating it plays a critical role in ligand recognition (Figure 2A).

**Figure 2.**
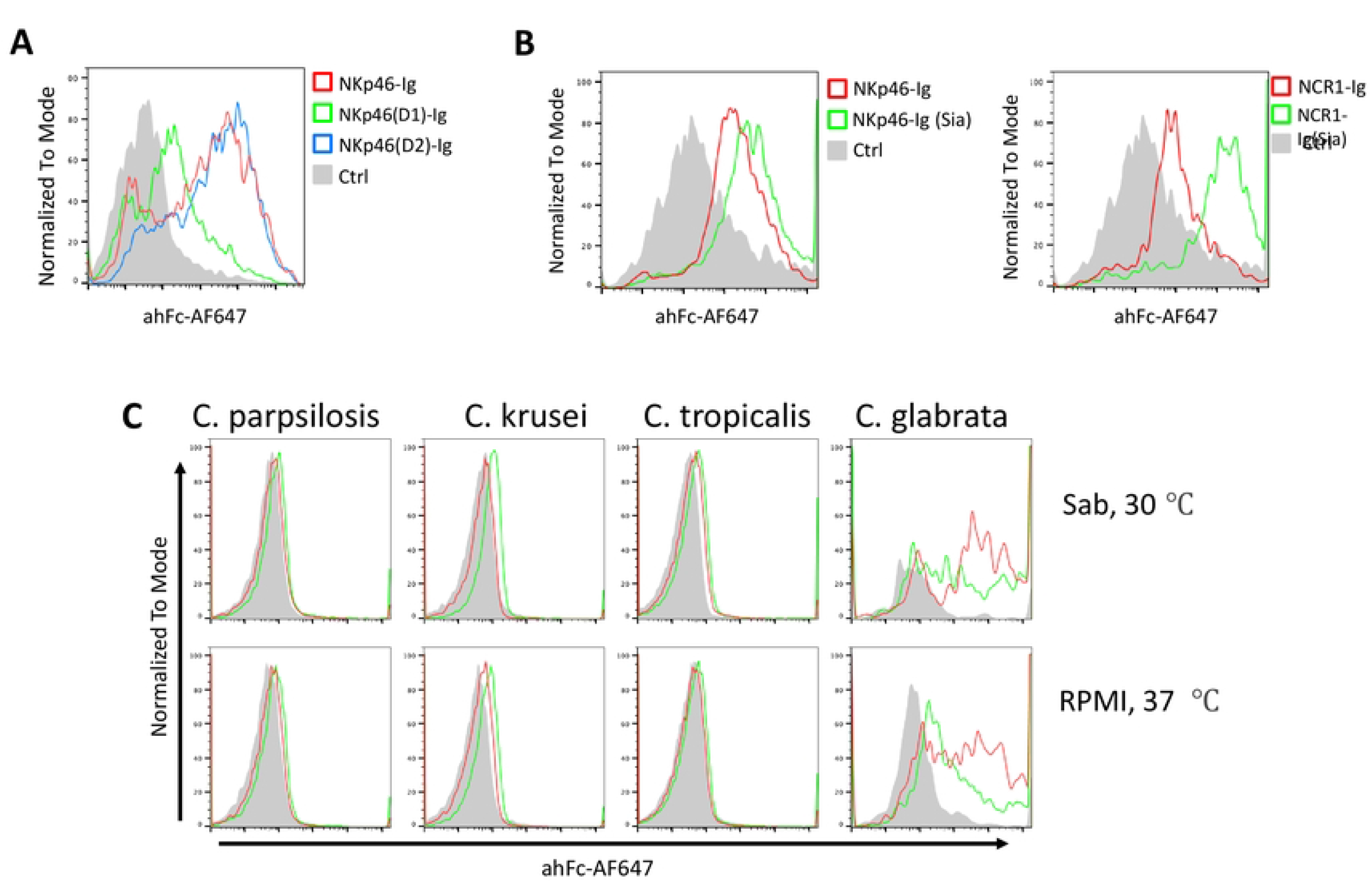
Binding properties of NKp46 to *C. albicans*. (A) Full-length NKp46-Ig and its individual D1-Ig and D2-Ig domains were used to stain hyphal *C. albicans*, followed by flow cytometry (FACS) analysis. Experiments were repeated twice. (B) Hyphal *C. albicans* was stained with NKp46-Ig (left) and NCR1-Ig (right), with or without prior sialidase (Sia) treatment, followed by FACS analysis. Experiments were repeated twice. (C) *C. parapsilosis, C. krusei, C. tropicalis*, and *C. glabrata* were cultured in Sabouraud medium at 30°C (upper) or in RPMI at 37°C (lower), then stained with NKp46-Ig and NCR1-Ig, followed by an anti-human Fc AF647-conjugated secondary antibody. Experiments were repeated twice.

Given the importance of sialic acid in other NKp46-ligand interactions[17,18], we treated NKp46-Ig and NCR1-Ig with sialidase prior to staining. Unexpectedly, sialidase treatment enhanced rather than diminished the binding of both NKp46-Ig (Figure 2B, left) and NCR1-Ig to *C. albicans* hyphae (Figure 2B, right), suggesting that sialic acid on NKp46 may partially mask its binding capacity in this context.

We next evaluated NKp46 and NCR1 binding to other *Candida* species under both yeast- and hyphae-inducing conditions. Consistent with previous findings, NKp46-Ig bound to *C. glabrata* under yeast-permissive conditions (30°C, Sabouraud medium, Figure 2C, upper), but not to other species[14]. Hyphal induction (37°C, RPMI, Figure 2C, lower) did not enhance NKp46 binding to species other than *C. albicans* (Figure 2C). To try and identify the Hyphal ligand of NKp46 we stained several *C. albicans* mutants lacking major adhesins and observed no change in NKp46-Ig binding, suggesting that NKp46 does not recognize known surface adhesins (Supplementary Figure 1).

### Blocking NKp46 impairs NK cell-mediated responses to *C. albicans*

To assess the functional relevance of NKp46 binding to hyphal *C. albicans*, we performed NK cell degranulation and killing assays. For this we used the NKp46-Ig protein as a competitive inhibitor. Human NK cells were co-cultured with *C. albicans* in the presence or absence of NKp46-Ig fusion protein. Addition of NKp46-Ig significantly reduced CD107a expression on NK cells (Figure 3A) and impaired their ability to kill *C. albicans* (Figure 3B), indicating that NKp46-mediated recognition of hyphae contributes to effective antifungal activity.

**Figure 3.**
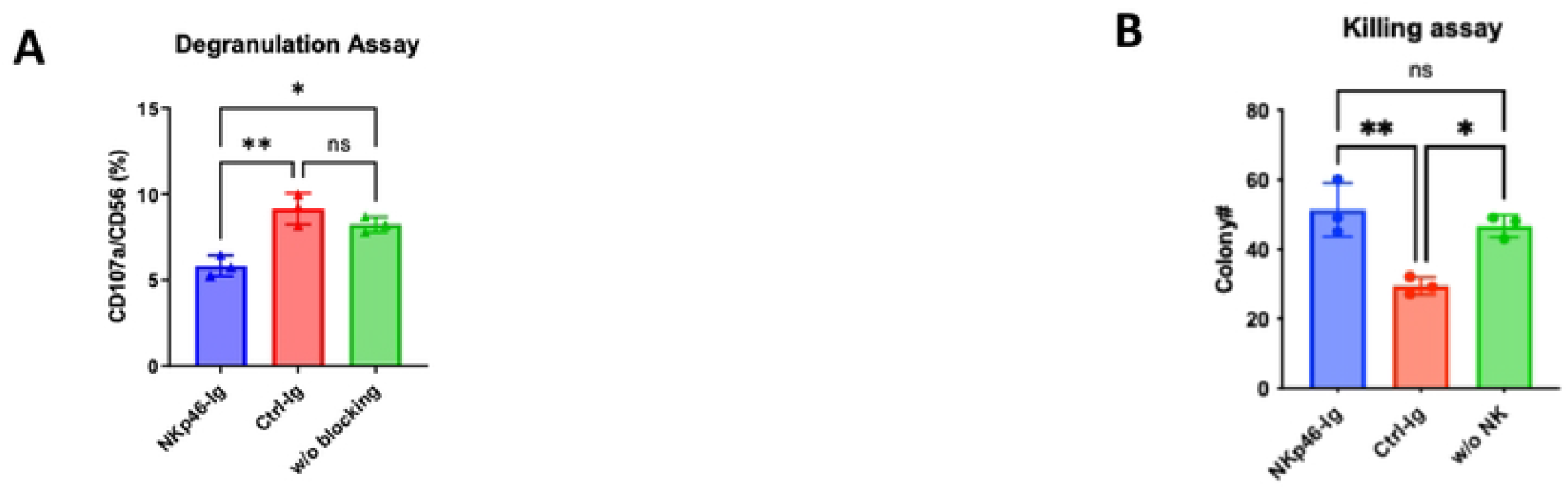
Blocking NKp46 reduces human NK cell-mediated killing of *C. albicans*. (A) Human NK cells were co-cultured with *C. albicans* for 4 hours with or without the indicated Ig proteins (X-axis) and stained with anti-CD56-PE and anti-CD107a-APC antibodies. The percentage of CD107a^+^ expression on CD56^+^ NK cells was analyzed by flow cytometry. Experiments were repeated twice. Statistical analysis was performed using ordinary one-way ANOVA followed by Tukey’s multiple comparisons test. Statistical significance is indicated as follows: *P* < 0.05 (*), *P* < 0.01 (**). (B) *C. albicans* and human NK cells were co-cultured overnight, and the number of viable *C. albicans* cells was quantified by serial dilution and plating on agar. Statistical analysis was performed using ordinary one-way ANOVA followed by Tukey’s multiple comparisons test. Statistical significance is indicated as follows: *P* < 0.05 (*), *P* < 0.01 (**).

### NCR1-deficient mice exhibit increased susceptibility to systemic *C. albicans* infection

To determine the in vivo relevance of NKp46 in host defense against *C. albicans*, we intravenously injected male and female wild-type and NCR1 knockout mice with *C. albicans* and monitored survival and body weight. NCR1-deficient mice displayed significantly increased mortality and more rapid weight loss compared to wild-type controls, in both male and female mice (Figure 4A, 4B).

**Figure 4.**
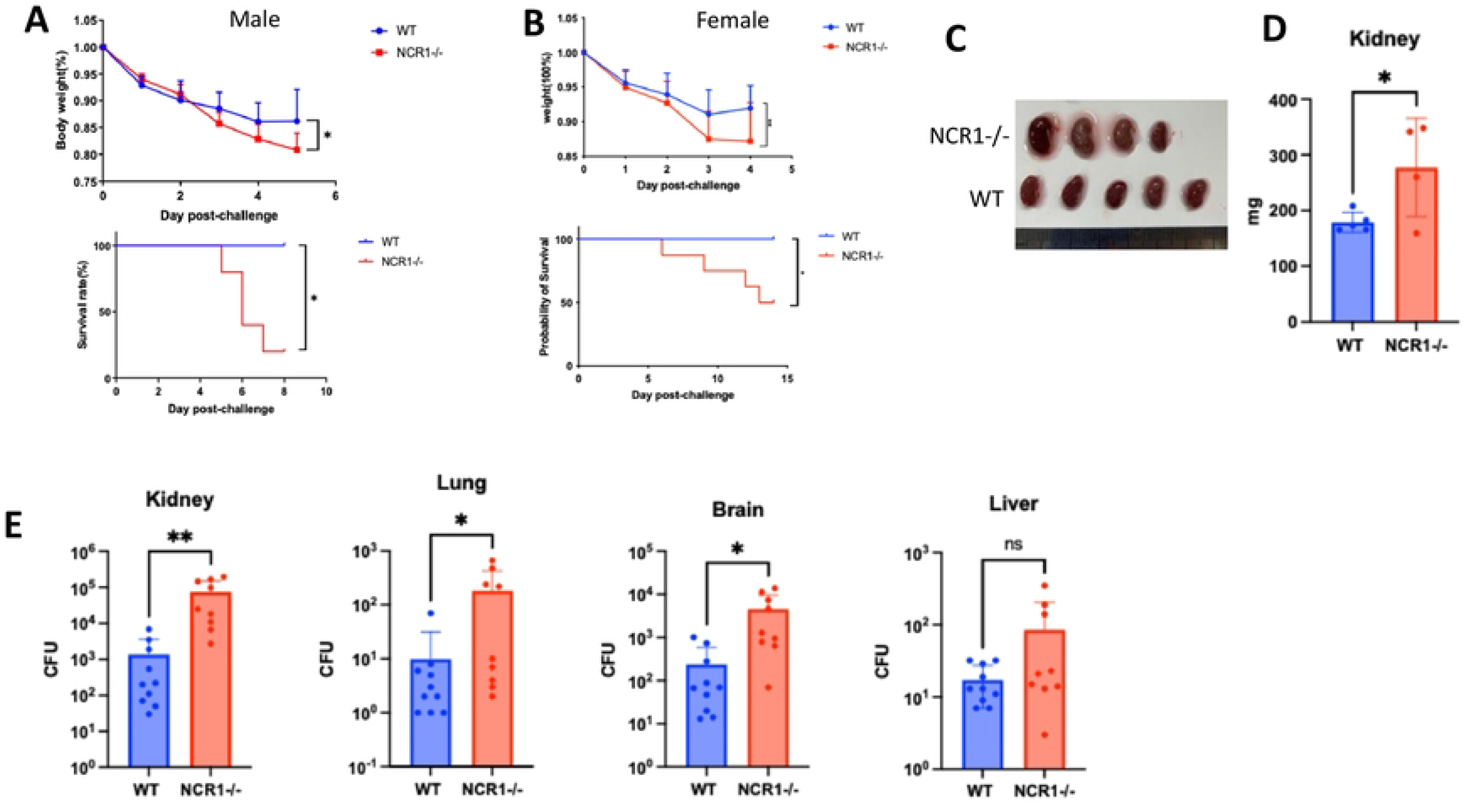
NCR1 knockout (KO) mice exhibit reduced resistance to *C. albicans* infection. (A) Wild-type (WT) (n=5) and NCR1 KO (n=5) male mice were injected intravenously with *C. albicans* (0.25M). Body weight was recorded daily. Survival rates of *C. albicans*-infected WT and NCR1 KO mice were assessed. The experiment was repeated twice. Statistical significance for weight differences was analyzed using two-way ANOVA with Šídák’s multiple comparisons test. Survival curves were analyzed using the Kaplan-Meier method, and statistical significance was determined by the Log-rank (Mantel-Cox) test. Statistical significance is indicated as follows: *P* < 0.05 (*), *P* < 0.01 (**). (B) Wild-type (WT) (n=10) and NCR1 KO (n=8) female mice were injected intravenously with *C. albicans* (0.25M). Body weight was recorded daily. Survival rates of *C. albicans*-infected WT and NCR1 KO mice were assessed. Statistical significance for weight differences was analyzed using two-way ANOVA with Šídák’s multiple comparisons test. Survival curves were analyzed using the Kaplan-Meier method, and statistical significance was determined by the Log-rank (Mantel-Cox) test. Statistical significance is indicated as follows: *P* < 0.05 (*), *P* < 0.01 (**). (C) Gross pathology of kidneys 2 days post-infection. (D) Kidney mass from WT and NCR1 KO mice was measured 2 days post-infection. Statistical significance was determined using an unpaired two-tailed Student’s t-test. Statistical significance is indicated as follows: *P* < 0.05 (*), *P* < 0.01 (**). (E) Fungal burden in the kidney, lung, brain, and liver 2 days post-*C. albicans* infection. Organs were homogenized and serially diluted (10×, 100×, 1000×). Diluted samples were plated on Sabouraud agar and incubated overnight at 30°C. Colonies were counted, and fungal burden was quantified. Experiments were repeated twice. Statistical significance was determined using an unpaired two-tailed Student’s *t*-test (*P* < 0.05 was considered statistically significant). Statistical significance is indicated as follows: *P* < 0.05 (*), *P* < 0.01 (**).

At day 2 post-infection, gross examination revealed enlarged kidneys in NCR1-deficient mice (Figure 4C), with significantly increased kidney mass compared to wild-type mice (Figure 4D). Fungal burden was also significantly elevated in multiple organs, including the kidney, brain, and lung (Figure 4E), suggesting compromised fungal clearance in the absence of NCR1.

We also examined the proportions of CD4^+^ T cells, CD8^+^ T cells, and NK cells in mice before and after *C. albicans* injection. NCR1^−^/^−^ mice exhibited a higher percentage of splenic NK cells, likely due to a compensatory expansion in the absence of NCR1 (Supplementary Figure 2). However, this increased NK cell frequency did not lead to enhanced fungal clearance, further indicating that NCR1 plays a critical functional role in the immune response against *C. albicans* infection.

NK cells have been reported to play an important role in *C. albicans* skin infection models involving deep dermal injections, where NK cell-depleted mice exhibit reduced ulceration compared to wild-type controls[19]. Based on these findings, we also investigated whether NCR1 contributes to this process. In contrast, NCR1 deficiency did not impact outcome in a cutaneous *C. albicans* infection model (Supplementary Figure 3), indicating that NKp46 plays a context-dependent role in host defense, particularly in systemic candidiasis.

## Discussion

Natural killer (NK) cells play a critical role in antifungal immunity, including against *C. albicans* and other pathogenic fungi [7,13,15]. Several NK cell receptors—such as TIGIT and NKG2D—have been shown to modulate NK cell responses to *C. albicans*.

The ability to distinguish between yeast and hyphal forms of *C. albicans* is crucial for the immune system. Hyphae are considered the more invasive morphology and are essential for mucosal penetration[4,20]. It has been previously shown that some immune molecules, such as Dectin-2 and mannose-binding lectin (MBL), preferentially bind the hyphal form of *C. albicans*, potentially enabling the immune system to target the most pathogenic form of the fungus[21,22]. Neutrophils, for example, exhibit enhanced phagocytosis and killing of hyphae compared to yeast, suggesting that early recognition of hyphae is a critical component of the antifungal response[23].

Our data show that NKp46 is another pattern recognition receptor capable of selectively identifying *C. albicans* hyphae. This interaction is mediated through the D2 domain of NKp46 and is independent of sialic acid, which is commonly involved in ligand interactions with NKp46 in other contexts such as viral and tumor recognition[17]. Interestingly, removal of sialic acid from NKp46 increased its binding to hyphae, suggesting a possible steric hindrance by sialic acid residues. These findings raise the possibility that NKp46 recognizes a non-sialylated fungal ligand, potentially a carbohydrate or glycoprotein component of the hyphal cell wall. Functionally, NKp46 binding to *C. albicans* hyphae is important for effective NK cell activation, as blocking this interaction significantly impairs NK cell degranulation and fungal killing. In vivo, NCR1-deficient mice were more susceptible to systemic *C. albicans* infection, exhibiting greater mortality, increased fungal burden, and severe kidney pathology. These data underscore the importance of NKp46 in mediating early immune responses against disseminated candidiasis.

Although the precise ligand of NKp46 on *C. albicans* remains to be identified, our attempts to screen knockout strains for loss of binding and to immunoprecipitate potential ligands from the fungal cell wall were inconclusive. This could be due to the fact that many knockout strains affecting hyphal formation also alter cell wall architecture, confounding the identification of specific ligands[24,25]. Nevertheless, the binding pattern and biochemical characteristics of this interaction suggest that the NKp46 ligand may be a hyphae-specific polysaccharide or an uncharacterized glycoprotein exposed during morphogenesis.

In summary, our study demonstrates that NKp46 is a functional receptor for *C. albicans* hyphae, contributing to NK cell-mediated antifungal immunity. This finding not only expands our understanding of NK cell recognition mechanisms in fungal infection but also highlights the importance of immune discrimination between commensal and pathogenic fungal forms. Targeting NKp46-ligand interactions may offer new strategies for enhancing antifungal immunity, particularly in immunocompromised patients at risk for invasive candidiasis.

## Methods

### Mice

Ncr1^gfp/gfp^ (KO) and C57BL/6 mice (6–8 weeks old; Envigo, Israel) were used in this study. All animals were specific pathogen-free (SPF), experimentally naïve, and group-housed. Littermates were randomly assigned to experimental groups. All procedures were performed in the SPF animal facility of the Hebrew University-Hadassah Medical School (Ein Kerem, Jerusalem), following the Declaration of Helsinki and institutional ethical guidelines.

### Human NK Cell Isolation and Culture

Human NK cells were isolated from peripheral blood mononuclear cells (PBMCs) of healthy donors using the EasySep™ Human NK Cell Enrichment Kit (STEMCELL Technologies). NK cells (5 × 10^4^/well) were co-cultured in U-bottom 96-well plates with irradiated (6000 RAD) PBMCs from two independent donors (5 × 10^4^/well per donor) and RPMI-8866 cells (5 × 10^3^/well). Cells were maintained in RPMI 1640 supplemented with 10% human serum (Sigma), 1 mM sodium pyruvate, 2 mM glutamine, non-essential amino acids, 100 U/mL penicillin, 0.1 mg/mL streptomycin (all from Biological Industries), 500 U/mL recombinant human IL-2 (PeproTech), and 20 μg/mL phytohemagglutinin (PHA; Sigma). Cultures were maintained at 37°C in 5% CO_2_. NK cell identity was verified by flow cytometry using anti-human CD56-PE and CD3-FITC antibodies (BioLegend).

### Microbial Strains

The fungal strains used included *Candida albicans* SC5314, *Candida glabrata* BG2, *Candida parapsilosis, Candida krusei*, and a panel of *C. albicans* deletion mutants (als1–als7Δ/Δ, als9Δ/Δ, ECE1Δ/Δ, HWP2Δ/Δ, HYR1Δ/Δ, RBT1Δ/Δ), all derived from SC5314.

Fungi were stored at −80 °C in glycerol stocks and streaked onto Sabouraud Dextrose Agar (SDA; Sigma) for up to 4 weeks. For experiments, colonies were cultured overnight in Sabouraud broth at 30°C with shaking, diluted 1:50 in fresh broth, and grown for an additional 2–4 h. For hyphal induction, cells were further diluted 1:20 into RPMI-1640 medium and incubated as above.

### Fusion Proteins

The extracellular domains of fusion proteins were cloned into a mammalian expression vector (pIRESpuro3) encoding a mutated human IgG1 Fc domain. Fusion proteins included NKp46-Ig, NKp46-D1-Ig, NKp46-D2-Ig, NTBA-Ig (negative control), and mouse NCR1-Ig, as described previously. Desialylation of NKp46-Ig was performed using neuraminidase beads (Sigma) as previously reported.

### Flow Cytometry and Imaging Flow Cytometry

Mammalian and fungal cells were prepared as described above, washed 3 times in ice-cold PBS (515 g for mammalian, 3000 g for fungi, 5 min at 4°C), and seeded into U-bottom 96-well plates (5–10 × 10^4^ cells/well). For blocking, cells were incubated with 2.5 μg/well of antibody for 30 min on ice. Cells were stained with 0.25 μg/well primary antibodies or 0.5–5 μg/well fusion proteins in FACS buffer (PBS, 0.05% BSA, 0.05% NaN_3_) for 1 h on ice. If necessary, cells were then stained with 0.75 μg/well fluorophore-conjugated secondary antibodies (AlexaFluor647 or APC; Jackson ImmunoResearch) for 30–45 min. Data were acquired on a CytoFlex (Beckman-Coulter) or ImageStream®X Mark II and analyzed using FCS Express software (De Novo Software).

### Fungal Killing and Degranulation Assays

Effector cells were blocked with NTBA-Ig or anti-NKp46-Ig (5 μg/well) for 1 h on ice. Cells were co-cultured with fungal targets (5 × 10^4^ effector cells and 1 × 10^3^ fungi/well) in RPMI-1640 medium for 12–14 h at 37 °C in 5% CO_2_. After incubation, cultures were serially diluted in PBS and plated on SDA. Plates were incubated at 30°C for 24–48 h, and colony-forming units (CFUs) were counted.

For degranulation assays, NK cells were co-cultured with fungal cells as described above. Anti-CD45-PE and anti-CD107a-APC antibodies (0.5 μl each per well) were added to the culture at the beginning of co-incubation. After 2 hours, cells were washed twice with cold FACS buffer and analyzed using a CytoFlex flow cytometer (Beckman-Coulter).

### Mouse Infection Model

On the day of infection, *C. albicans* cells were washed 3 times in PBS and kept on ice. Mice were intravenously injected with 2.5 × 10^5^ CFU in 100 μL PBS via the tail vein.

#### Fungal Burden

At 48 h post-infection, mice were euthanized and organs harvested. Organs were homogenized, filtered through a 70 μm strainer, serially diluted in PBS, and plated **on SDA. CFUs were counted after 48 h of incubation at 30 °C.**

#### Survival Analysis

Mice were monitored daily. Animals were euthanized if body weight dropped below 80% of baseline or if severe clinical signs developed.

## Acknowledgments

Ofer Mandelboim is supported by: The Israel Science Foundation grant 3042/22, grant 442/18 and grant 619/23. The German-Israeli Foundation for Scientific Research and Development Grant 1412-414.13/2017. The Israel Innovation Authority grant 75934. The ICRF professorship. The Yissum grant 220318 The MOST-DKFZ grant 3-14931 and grant 3 14930

## Figure legends

**Supplementary Figure 1. Major adhesion molecules of *C. albicans* are not NKp46 ligands.**

(A) *C. albicans* strains lacking key adhesion molecules were stained with NKp46-Ig, followed by an anti-human Fc AF647-conjugated secondary antibody. Experiments were repeated twice.

**Supplementary Figure 2. NCR1-deficient (NCR1^−^/^−^) mice exhibit an increased frequency of NK cells in the spleen compared to wild-type controls.**

(A) The proportions of CD4 T cells, CD8 T cells, and NK cells in the spleen of mice without *C. albicans* infection were analyzed by flow cytometry. (B) The proportions of CD4 T cells, CD8 T cells, and NK cells in the spleen of mice after *C. albicans* injection were analyzed by flow cytometry.Statistical significance was determined using an unpaired two-tailed Student’s *t*-test (*P* < 0.05 was considered statistically significant). Statistical significance is indicated as follows: *P* < 0.05 (*), *P* < 0.01 (**).

**Supplementary Figure 3. No significant difference between WT and NCR1 KO mice in a skin infection model.**

Percentage of WT and NCR1 KO mice developing ulceration following *C. albicans* administration. Experiments were repeated twice. Survival analysis was performed using the Kaplan-Meier method, and statistical significance was determined by the Log-rank (Mantel-Cox) test. Statistical significance is indicated as follows: *P* < 0.05 (*), *P* < 0.01 (**).

